# A combination of Mechanical, Chemical, and Thermal Pretreatments of Agricultural Feedstocks Enhances Biomethane Yields in Advanced Anaerobic Digestion

**DOI:** 10.1101/2025.04.02.646888

**Authors:** Nichola Austen, Helen Theaker, Nicholas A. Tenci, Alan Beesley, Ian P. Thompson

**Affiliations:** Department of Engineering Science, University of Oxford, Oxford, OX1 3SR, UK; Carl R. Woese Institute for Genomic Biology, University of Illinois at Urbana-Champaign, IL, 61801, USA; Center for Advanced Bioenergy and Bioproducts Innovation, University of Illinois at Urbana-Champaign, IL, 61801, USA; ALPS Ecoscience Ltd, 15 The Metro Centre, Toutley Road, Wokingham, Berkshire, RG41 1QW, UK

**Keywords:** Bioenergy crops, Anaerobic digestion, Biogas yield, Feedstock Treatment, Bioeconomy, UK net zero, Hydrolysis

## Abstract

Multi-step Advanced Anaerobic Digestion (AAD) pretreatment of feedstocks increases biogas yields compared to non-pretreated feedstocks and is key to the processing of recalcitrant lignocellulosic feedstock to make commercial biogas production more economically viable. Here, we present several low energy and eco-friendly pretreatments to commercially relevant lignocellulosic feedstocks (rye and maize), to increase biomethane yields. In this study the impact of two heating treatments, 55 °C and at 80 °C, the addition of a bio-organic catalyst (BOC), and mechanical particle size reduction by cavitation were investigated. For both feedstocks, thermal pretreatment significantly increased both solubility and enhanced biogas yield (8.6 – 136.6%), with maize responding better to a temperature of 55 °C (136.6% increase) and rye to 80 °C (62% increase). The BOC addition enhanced the rye yield (14%) but decreased from maize (4%), and cavitation enhanced the Biochemical Methane Potential (BMP) of rye (38.7%) but had an inhibitory effect on maize (10.6%). The results of this multi-process study demonstrate the efficacy of low energy pretreatments for lignocellulosic material that can be applied to existing AD plants.

## 1. Introduction

Improvement of renewable and sustainable energy generation is urgently needed to match and replace the current consumption of fossil fuels to help to mitigate the climate crisis. Anaerobic digestion (AD) is a well-established technology that can be used as part of a blended clean energy approach. Traditionally, AD has been applied more in rural areas in countries such as China and India [1, 2], with AD in China accounting for > 10% of energy generation in its rural areas [1]. In Europe and the USA, there has been significant development of AD as an energy source over the last 20 years, with over 9000 plants established in Germany alone [3], and over 2300 in the USA [3]. In the UK specifically, as of 2023 2.2% of agricultural land is used for growing bioenergy crops, with most of this being used to grow maize for AD [4]. Traditionally, AD feedstocks have been waste products from agriculture, food, and animal and human waste [5]. Often, the plant biomass used is rich in lignocellulosic material, which is difficult to break down by the microbial communities and leads to long lag phases in biogas generation [6]. In AD, hydrolysis is the rate limiting step [7, 8], increasing the time to biogas production through the slow breakdown of solid, lignocellulosic material. Preprocessing these recalcitrant feedstocks provides a more readily available soluble material and can significantly improve conversion efficiencies, reduce reactor hydraulic retention times, and ultimately increase methane yields [9-11].

An increase in methane yield and, importantly, increases in additional high value volatile fatty acids will be highly relevant to the renewable energy and biorefinery industries, existing biogas plants [12, 13], and sustainable chemical production sectors, to decrease dependence on petrochemicals. Commercialisation of novel pre-processing systems has strong industrial drivers, enabling more biogas to be released from the same amount of feedstock, reducing the digestion volume required and production costs, and promoting the efficient use of land resources for the plant footprint and for growing feedstocks [14, 15]. The process changes that are required can be retrofitted to existing infrastructure with minimal additional investment.

Advanced anaerobic digestion (AAD) is a term used to characterize AD plants with additional processing capabilities and can be used to describe any AD processes where extra steps are added, specifically before the feedstock reaches the main digester, to increase efficiency. ALPS Ecoscience have conducted prior studies on a range of pre-processing strategies and offer these commercially, including mechanical particle size reduction by cavitation, addition of a heating stage, acid hydrolysis and two-stage digestion incorporating dark fermentation. The continuing economics of existing AD plants will rely on being able to run at a greater conversion efficiency, when subsidies such as Renewable Heat Incentive and Feed-In Tariffs expire after 15-20 years from commissioning, in 2026 [16]. In conjunction with cavitation, ALPS Ecoscience employs a novel chemical surfactant additive called Eco-Cat Bio-Organic Catalyst or BOC (Bio-Organic Catalyst Inc, Costa Mesa, USA). BOCs contain a fermentation supernatant (derived from plants and minerals), and surface modifying components which in water create ultra-fine micro-bubbles for accelerated chemical and biological reactions and higher conversion rates within water and solid waste treatments. The breaking of wastes (such as fats, oil, and greases (FOGs), food waste, and lignocellulosic materials in AD) into their constituent components create more digestible products for microorganisms [17]. When lignocellulosic bonds are broken, this additive binds to the isolated lignin, leaving cellulose free for digestion, thus boosting the organic nutrients available for digestion [18, 19]. In commercial operations, BOC has proven effective in maintaining the separation between molecules, ensuring that post cavitation, the digestion of the cellulosic material is not inhibited by lignin rebinding. This increases the degradation potential when compared to untreated material.

In this study the effect of thermal pretreatment in conjunction with both cavitation (mechanical pretreatment) and the addition of Eco-Cat (chemical pretreatment) on gas yields for the first time were systematically investigated. Laboratory-scale experiments were designed to investigate and validate a series of novel pretreatments, designed by ALPS Ecoscience on different feedstocks, to measure the potential improvements in biogas yields, shorter retention times, and improved efficiency of AD through a two-step process. A biomethane potential assay (BMP) was conducted on several different pretreatments of two feedstocks: maize and rye which were compared with the untreated feedstocks as a control. Maize and rye grass were chosen as feedstocks for the experiments since they are commercially important feedstocks for AD in the UK. From 2019 onwards, ∼ 75,000 Ha of agricultural land was used to cultivate maize for bioenergy [4]. Although the area of land used to cultivate rye in the UK has increased to around 50,000 Ha this year, the use of rye as an AD feedstock has declined [20], due to economic implications.

Biogas yield results with relevant soluble chemical oxygen demand (SCOD) and subsequent statistical analyses are presented, to quantify the improvement of biomethane generation.

## 2. Methods

### 2.1 Biomass and inoculum

Maize (*Zea mays*) and rye (*Secale cereale*) samples were collected from the Engie UK Ltd Energy biogas plant, which is operating at Frogmary Green Farm, in Somerset, England. The maize was sewn in early May 2023, and the rye was drilled as a cover crop in late September 2022. The maize was harvested at maturity in late September, and the rye was harvested in July 2023. Both crops were stored in a silage clamp prior to collection for the experiment. The AD sludge was collected from Cassington Anaerobic Digestion Facility, Oxfordshire UK (Severn Trent Green Power) and was stored under anaerobic conditions at 30 °C before use. Table 1 summarises the inoculum characteristics. The agricultural feedstocks were pretreated at ALPS Ecoscience test facility (Chedglow, UK) as follows: maize and rye were treated separately but under the same conditions. 10 kg of chopped (∼ 2cm), homogenised feedstock (maize and rye run separately) was added to a 220 L mixing tank with 33 L of water and mixed constantly for 30 minutes. A cavitation pump (6 m^3^/hr “Biobang”, Soldo Cavitators, Italy) was used and the feedstock and water mixture and pumped around the system 10 times.

**Table 1.** Bulk biomass characteristics (%) of substrate and inoculum used in the experiment. All data are measured in triplicate and given as means ± of standard deviation.

| Characteristic | Substrate |  |  |  |  |  |  |  |  |  |  | Inoculum |  |
| --- | --- | --- | --- | --- | --- | --- | --- | --- | --- | --- | --- | --- | --- |
|  | Maize<sub>2</sub> | Rye<sub>2</sub> | Maize+<br>CAV<sub>1</sub> | Rye+<br>CAV<sup>2</sup> | Maize+<br>BOC+<br>CAV<sub>2</sub> | Rye+<br>BOC+<br>CAV<sub>2</sub> | BOC | Maize+<br>Heat<br>(80 °C)<sub>2</sub> | Rye+<br>Heat (80<br>°C)<sub>2</sub> | Maize+<br>Heat<br>(55 °C)<sub>1</sub> | Rye+<br>Heat<br>(55 °C)<sub>1</sub> | Sludge<br>1 | Sludge<br>2 |
| TS/FW (%) | 30.10<br>$\pm 1.85$ | 28.84<br>$\pm 0.33$ | 2.32<br>$\pm 0.24$ | 3.16<br>$\pm 0.03$ | 2.37<br>$\pm 0.10$ | 2.48<br>$\pm 0.02$ | 11.49<br>$\pm 0.10$ | 2.63<br>$\pm 0.27$ | 6.80<br>$\pm 1.14$ | 1.72<br>$\pm 0.33$ | 1.69<br>$\pm 0.14$ | 4.01<br>$\pm 0.02$ | 1.94<br>$\pm 0.02$ |
| VS/TS (%) | 97.15<br>$\pm 0.22$ | 92.82<br>$\pm 2.92$ | 95.09 ±<br>1.04 | 56.42<br>$\pm 1.21$ | 92.99<br>$\pm 1.69$ | 76.02<br>$\pm 0.96$ | 94.75<br>$\pm 0.96$ | 92.49<br>$\pm 1.04$ | 87.31<br>$\pm 1.91$ | 97.29<br>$\pm 1.14$ | 83.68<br>$\pm 1.04$ | 63.57<br>$\pm 0.34$ | 52.73<br>$\pm 2.51$ |
| Ash/TS (%) | 2.85<br>$\pm 0.21$ | 7.18<br>$\pm 2.92$ | 4.91<br>$\pm 1.00$ | 43.58<br>$\pm 1.20$ | 7.01<br>$\pm 1.69$ | 23.98<br>$\pm 0.96$ | 5.25<br>$\pm 0.95$ | 7.51<br>$\pm 1.00$ | 12.69<br>$\pm 1.91$ | 2.71<br>$\pm 1.00$ | 16.32<br>$\pm 1.00$ | 36.43<br>$\pm 0.34$ | 47.27<br>$\pm 2.51$ |

\* TS – total solids, FW – fresh weight, VS – volatile solids, <sup>1</sup>= AD sludge 1, <sup>2</sup> = AD sludge 2

Once a viable sample had been drawn off for testing (5 L), Eco-Cat bio-organic catalyst (BOC) (Nijhuis Industries, Wokingham, UK) was added in a ratio of 1 mL/kg of Organic Dry Matter and left to mix in the tank for 30 minutes. A 5 L sample of the pretreated maize and water mixture was also drawn off for testing. The same procedure was undertaken for the rye feedstock. Further experimental test samples were made in the laboratory (Oxford, UK) and all pretreatments are described in full in Supplementary Table 1.

Following an initial determination of bulk biomass parameters, the industrial scale samples (which included the following treatments for both maize and rye feedstocks: cavitated and cavitated with BOC) were determined to be too dilute for the laboratory scale experiment and were concentrated down in a drying oven to∼ 50% of their original volume (see Supplementary Table 1 for details) at a temperature of 80 °C for 48 hours. As the samples collected from ALPS were subject to heat to concentrate them, additional treatments were included to investigate the effects of thermal pretreatment on feedstocks that were not subject to cavitation. At the same ratio of feedstock to water as the industrial samples, two different heat treatments at 55 °C and 80 °C for 48 hours were prepared in the laboratory for both maize and rye feedstocks in 5 L schott bottles. Further pretreatments were the untreated feedstocks only, both maize and rye with added BOC only, and a thermal pretreatment of 80 °C with added BOC was prepared for each of the feedstocks. Due to issues with sample concentration, some pretreatments were run separately with a different batch of sludge. The following pretreatments were run with AD sludge 1 as an inoculum: maize + cavitation, maize + heat (55 °C), rye + heat (55 °C). All other treatments were run with AD sludge 2 as an inoculum. The differences in sludge biomass characteristics are detailed in Table 1.

### 2.2 Anaerobic batch assays

The biochemical methane potential experiment was conducted as a batch assay in 250 mL vials with a 150 mL working volume. A culture medium of macronutrients, micronutrients, minerals, vitamins and buffers (with the exclusion of sodium sulphide) was prepared according to the method of Angelidaki & Sanders [21] to support microorganism growth. The ratio of substrate to inoculum was set at 1:1 (w/w) ratio on a volatile solid (VS) basis. The mastermix pH was adjusted to 8 prior to the start of the experiment. The master mix, consisting of medium solution, deionized water, and the inoculum was added to the vials with the substrate (see Table 1 for substrate characteristics) which were sealed with rubber septa and aluminium crimp caps and were inverted to mix. Both a blank control (inoculum only) and positive control (microcrystalline cellulose) were included in the experiment with all treatments run in triplicate. All conditions were incubated at 37 °C with constant agitation of 125 RPM. The experiment was terminated on day 60.

### 2.3 Analytical methods

Characterisation of both the bulk feedstock and the AD sludge parameters (both Total and Volatile Solids (TS/VS)) were measured according to standard methods previously described in APHA [22].

Biogas production throughout the experiment was measured with a eudiometer, with measurements corrected for both standard temperature (273.15 K) and pressure (101.3 kPa) as previously described by Richards *et al*., [23]. At each time point, liquid samples of 600 µl were extracted from each vial using a syringe with the pH taken immediately after sample collection. The samples were then frozen at -80 °C for measurement of soluble chemical oxygen demand (SCOD).

The methane composition of the biogas was determined by Gas Chromatography Flame Ionisation Detection (GC-FID) analysis (Shimadzu GC-2010, Japan). The column used was a 30m x 0.53 mm x 20 µm HP-PLOT U capillary column. The temperatures of the detector, injector, and oven were 250, 180, and 30 °C, respectively. Helium was used as the carrier gas.

The liquid samples were centrifuged at 20, 000 x g for 10 mins where 200 µl was used for SCOD analysis. SCOD was determined for the initial time points (time point zero) of each of the samples. The supernatant was analysed using a LCI400 COD cuvette test and DR2800 spectrophotometer (Hach Lange).

### 2.4 Statistical analyses and modelling

The experimental data presented here were analysed in triplicate, with the mean ± standard deviation presented. All data analyses were conducted in Graphpad Prism 10.2.3 [24].

Biomethane yields were calculated as the sum of both the substrate and inoculum VS. Biomethane yield data were fitted using the following Modified Gompertz kinetic model, as previously demonstrated by Lueangwattanapong et al. [25]:

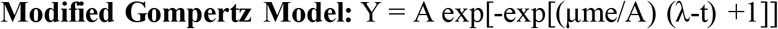

Where, Y = cumulative methane yield at time ‘t’, A = maximum theoretical methane yield, t = time, λ = lag phase, μm = maximum methane production rate. This model was fitted in GraphPad Prism 10.2.3 to the methane yield data for each treatment to produce the above kinetic parameters.

Statistical analyses were conducted using Graphpad Prism 10.2.3. One way analysis of variance (ANOVA) tests with subsequent *post-hoc* Tukey HSD tests (*p = 0.05*) were performed on both the SCOD data values, and the end time point value of the cumulative biomethane curves. The full results are in Supplementary Table 2.

## 3. Results

### 3.1 Characterisation of two agricultural feedstocks, pretreatments, and inoculum

The data in Table 1 detail the biomass characterization of the two agricultural feedstocks, the various pretreatments applied to them, and two commercial AD sludges used as the inocula for the experiment. The total solids (TS) as a proportion of fresh weight (FW) (TS/FW%) of the AD sludge 1 used (4.01 ± 0.02%) agree with previous studies using the same inoculum source [25, 26]. However, the AD sludge 2 collected had a lower TS (1.94 ± 0.02%) when compared to the literature. Volatile solid (VS) content of the TS data for AD sludge 1 was also consistent with previous studies, with batch one containing 63.57 ± 0.34%. Again, AD sludge 2 contained a lower VS percentage of TS in comparison to AD sludge 1 (52.73 ± 2.51%). Both maize and rye had similar amounts of TS/FM (30.1 ± 1.85 % and 28.84 ± 0.33% respectively). This contrasts with the pretreatments, which had significantly lower levels of TS/FW% in comparison to the raw feedstock. This ranged from a TS/FM of 1.69 ± 0.14% for rye + heat (55 °C) to 6.8 ± 1.14% for rye + heat (80 °C). In contrast, the BOC additive (which was added to the samples in liquid form) had a TS/FW of 11.49 ± 0.1%. All but four of the feedstocks and pretreatments had a VS/TS of over 92%, with both maize and maize + heat (55 °C) yielding the highest percentages of VS/TS, at 97.15 ± 0.22% and 97.29 ± 1.14% respectively. The pretreatment maize + cavitation had the lowest VS/TS at 56.42 ± 1.21%. Both rye + heat treatments (55 °C and 80 °C) had a similar TS/FM of 83.68 ± 1.04% and 87.31 ± 1.91% respectively.

### 3.2 Increases in solubility of two agricultural feedstocks due to novel pretreatments

To observe the potential differences between the raw agricultural feedstocks and the pretreatments, a measurement of solubility at time point zero of the biomethane potential (BMP) experiment was taken. This measurement gives an indication of the total organic material (g/L) that could be hydrolysed by the bacterial communities during the first stage of the AD process, demonstrated in Figure 1A and B. Both figures showed an increase in solubility when the two feedstocks were subjected to heat at varying temperatures when compared to other pretreatments. There was greater variability in solubility between treatments in the maize samples than there were both between the rye samples, and between the two different feedstocks.

**Figure 1.**
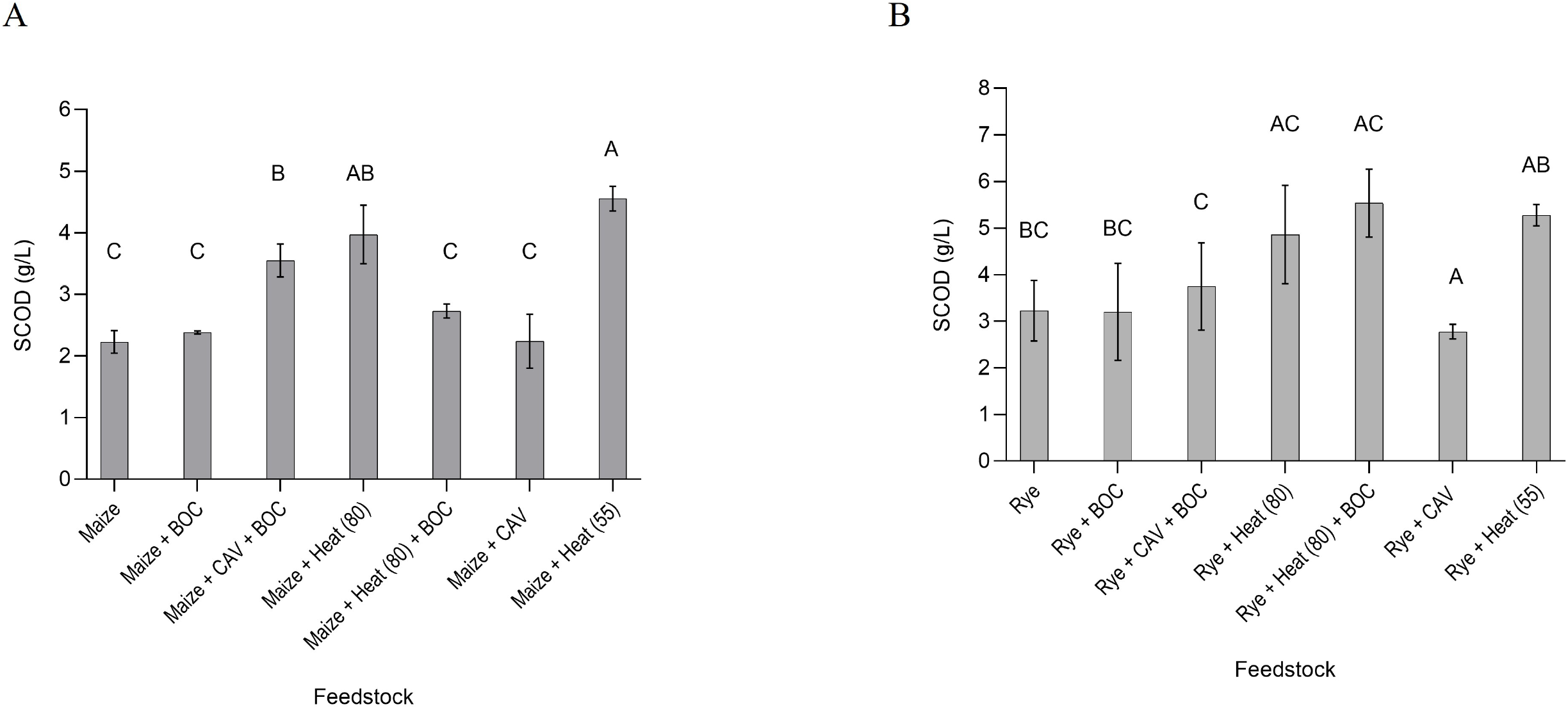
Soluble chemical oxygen demand (SCOD) (g/L) of each of the treatments for two pretreated agricultural feedstocks. A) Maize, and various pretreatments, B) Rye, and various pretreatments. Both graphs have been subject to a one-way ANOVA with *post-hoc* Tukey tests where the ANOVA summary is: *p =<0.0001* (maize) and *p =0.002105* (rye). Full details of one-way ANOVA results can be found in Supp. Table 2. All data presented as mean ±standard deviationof triplicate measurements. Results of Tukey’s HSD test of significant differences are shown on the graphs by letter.

Overall, the solubility of the rye feedstocks and pretreated samples were greater than the maize treatments. The average potential available soluble material of the untreated rye was 3.23 g/L compared to an average of 2.23 g/L for the untreated maize. Of all the pretreatments of the rye, the heated (80 °C) sample with added BOC had the greatest amount of potential soluble material at the start of the experiment. At 5.53 g/L, this was over a 60% increase in available material than the untreated rye. Both the other two rye heat treatments (rye + heat (80 °C) and rye + heat (55 °C)), yielded similar results, with an increase in theoretical solubility at > 46% and > 58%, respectively. In contrast, the maize heat pretreatments performed differently, with the maize + heat (55 °C) producing over 100% more Soluble Chemical Oxygen Demand (SCOD) than the untreated maize, closely followed by the maize + heat (80 °C) at 78% greater SCOD. In contrast to the rye, the maize + heat (80 °C) + BOC performed relatively poorly, with only a 22% increase in SCOD when compared with the untreated feedstock. In both maize and rye samples, both the cavitated samples (without added BOC) and the untreated feedstock (with and without added BOC) both had the lowest levels of soluble material available, when compared to the other treatments.

### 3.3 Biomethane yield of two different feedstocks

Although the rye feedstock and pretreatments revealed a greater amount of solubility at the outset, maize pretreatments demonstrated a higher biomethane yield (Figure 2 A and B).

**Figure 2.**
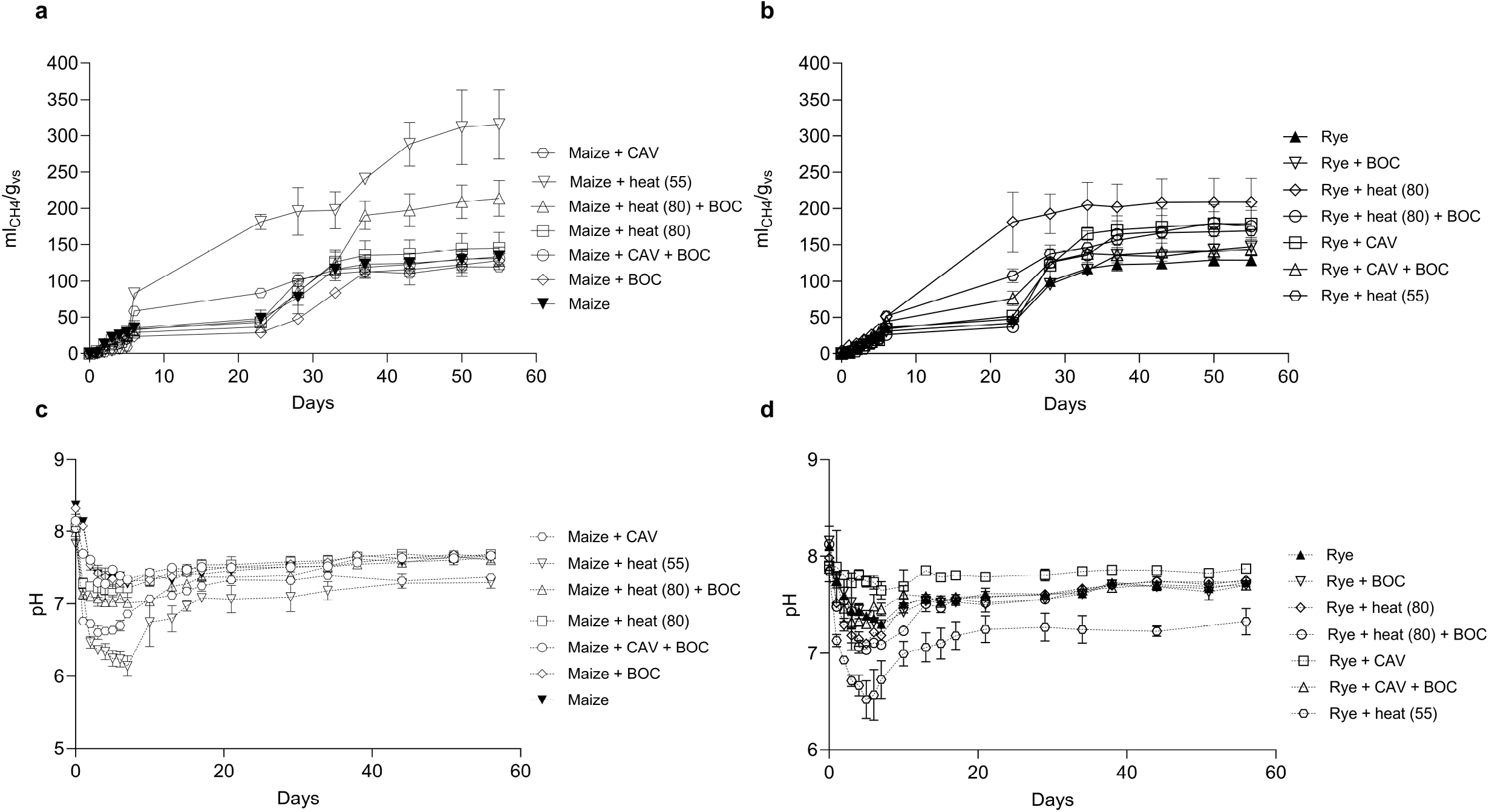
Total Biomethane yield and pH time courses for two agricultural feedstocks. A) maize CH_4_ yield (ml CH_4_g VS), B) Rye CH_4_ yield (ml CH_4_g VS), C) Maize pH, D) Rye pH. Yields are given on a g/VS basis. All data are presented as mean ± standard deviation of triplicate measurements.

Figure 2 C and D show the corresponding pH measurements across the time course experiment for both maize and rye treatments. The results of the BMP curves were fitted to a modified Gompertz model to interrogate the effects of different pretreatments of both maize and rye on theoretical maximum rates of production (µ_m_), theoretical total production (A), and the lag time (λ) of each experimental factor. The results of the non-linear regressions are shown in Figure 3A; maize control and pretreatments and Figure 3B rye control and pretreatments The results of the model parameters can be found in Table 2.

**Figure 3.**
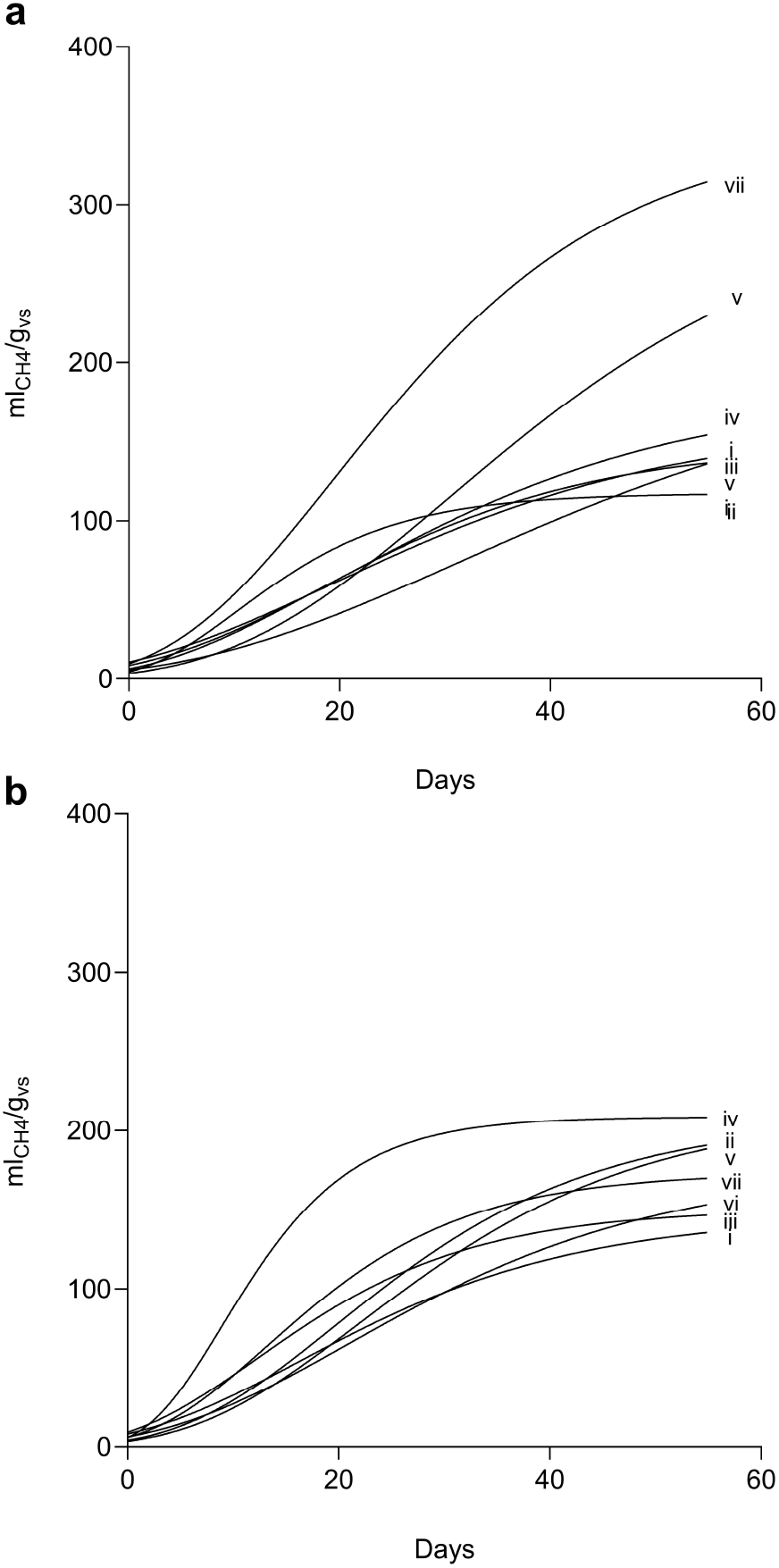
Modified Gompertz models of biomethane yields. The data represent models of best fir for the raw data shown in Figure 2 A & B. A) Maize – **i**. maize, **ii**. Maize + cav, **iii**. Maize + cav + BOC, **iv**. Maize + heat (80°C), **v**. maize + heat (80°C) + BOC, **vi**. Maize + BOC, **vii**. Maize + heat (55 °C). B) Rye - **i**. Rye, **ii**. Rye + cav, **iii**. Rye + cav + BOC, **iv**. Rye + heat (80°C), **v**. rye + heat (80°C) + BOC, **vi**. Rye + BOC, **vii**. Rye + heat (55 °C).

**Table 2.** Modelling parameters from modified Gompertz model of biomethane production from pretreated agricultural feedstocks.

| Treatment | Model parameters |  |  | Model R<sup>2</sup> |
| --- | --- | --- | --- | --- |
| | A (mL/ gVS) | $\mu_m$ (mL/ gVS/d) | $\lambda$ (d) | |
| Maize | 162.5 | 3.122 | 0.007664 | 0.9188 |
| Maize + CAV | 117.4 | 4.919 | 1.541 | 0.9404 |
| Maize + BOC | 197.9 | 2.969 | 6.397 | 0.9370 |
| Maize + CAV + BOC | 148.7 | 3.586 | 2.188 | 0.9512 |
| Maize + heat (55 °C) | 348.3 | 8.303 | 4.232 | 0.9558 |
| Maize + heat (80 °C) | 180.9 | 3.565 | 2.112 | 0.9189 |
| Maize + heat (80 °C)<br>+ BOC | 308.7 | 5.602 | 9.922 | 0.9346 |
| Rye | 146.7 | 3.531 | 1.037 | 0.9493 |
| Rye + CAV | 208 | 5.326 | 5.286 | 0.9467 |
| Rye + BOC | 176.2 | 3.742 | 3.529 | 0.9563 |
| Rye + CAV + BOC | 150.6 | 4.674 | 0.3323 | 0.9621 |
| Rye + heat (55 °C) | 173.6 | 5.889 | 2.405 | 0.9496 |
| Rye + heat (80 °C) | 208.7 | 10.89 | 1.943 | 0.9664 |
| Rye + heat (80 °C) + BOC | 212.6 | 5.036 | 6.452 | 0.9501 |

\* A – biomethane production potential, $\mu_m$ – max. production rate, $\lambda$ – lag phase.

The theoretical maximum biomethane production demonstrates higher yields from the maize than from the rye feedstock with a 64% increase in methane production in the maize + heat (55 °C) treatment. The theoretical highest maize pretreatment yield was 114% greater than the maize feedstock, and almost 200% greater than the yield for the maize + cavitation pretreatment. In contrast, there is a 45% yield increase between the rye feedstock and the highest yielding rye pretreatment (rye + heat (80 °C) + BOC) (Figure 3A, 3B and Table 2).

There were differences in the yields between maize pretreatments with added BOC, and those without. A significant yield increase of 71% is shown in Table 2 between the maize + heat (80 °C) and the maize + heat (80 °C) + BOC pretreatments. There was no significant increase however in the rye 80 °C heat treatments with added BOC from those without BOC (2%).

The addition of BOC to the cavitated rye samples appeared to have an inhibitory effect on the theoretical maximum yields of methane produced as shown in Table 2. There was a decrease in yield between the cavitated rye and rye + BOC of 15%, and a decrease in yield between the cavitated rye and rye + cavitation + BOC of 28%.

With the exception of maize + heat (80 °C), all heated pretreatments had higher theoretical daily maximum production rates, with rye + heat (80 °C) generating a daily rate of 10.89 mL/gVS/d, compared to daily rates of 3.122 mL/gVS/d and 3.531 mL/gVS/d for maize and rye respectively. There was significant variation (Table 2) in the length of the lag phase between the different feedstocks, with no clear trend of the effects of specific pretreatments on the degree of delay in methane production ranging from 0.007 – 9.92 days (maize – maize + heat (80 °C) + BOC).

There was significant variation in the differences of methane yield between pretreatments of the same feedstock, as well as differences between the two feedstocks and associated pretreatments as shown in Figs. 2 (A, B) and 3 (A, B). The differences in the total cumulative methane yield from the BMP assay are summarised in Figure 4 (A and B), with the final cumulative yield measurement provided. There was a significant difference (*p = < 0.0001*) between the maize feedstock and the heat (55 °C) pretreatment maize, with an approximate doubling of the final methane yield (Fig. 4A). The differences between the heat-treated samples (at both 80 °C and 55 °C) and the cavitated/cavitated with added BOC was also significant in maize samples (Fig. 4A). In contrast, the only significant differences between the methane yields of the rye samples was between the untreated residue (Fig. 4B), and the rye + heat (80 °C) treatment, as all other methane yields for the other pretreatments were comparable.

**Figure 4.**
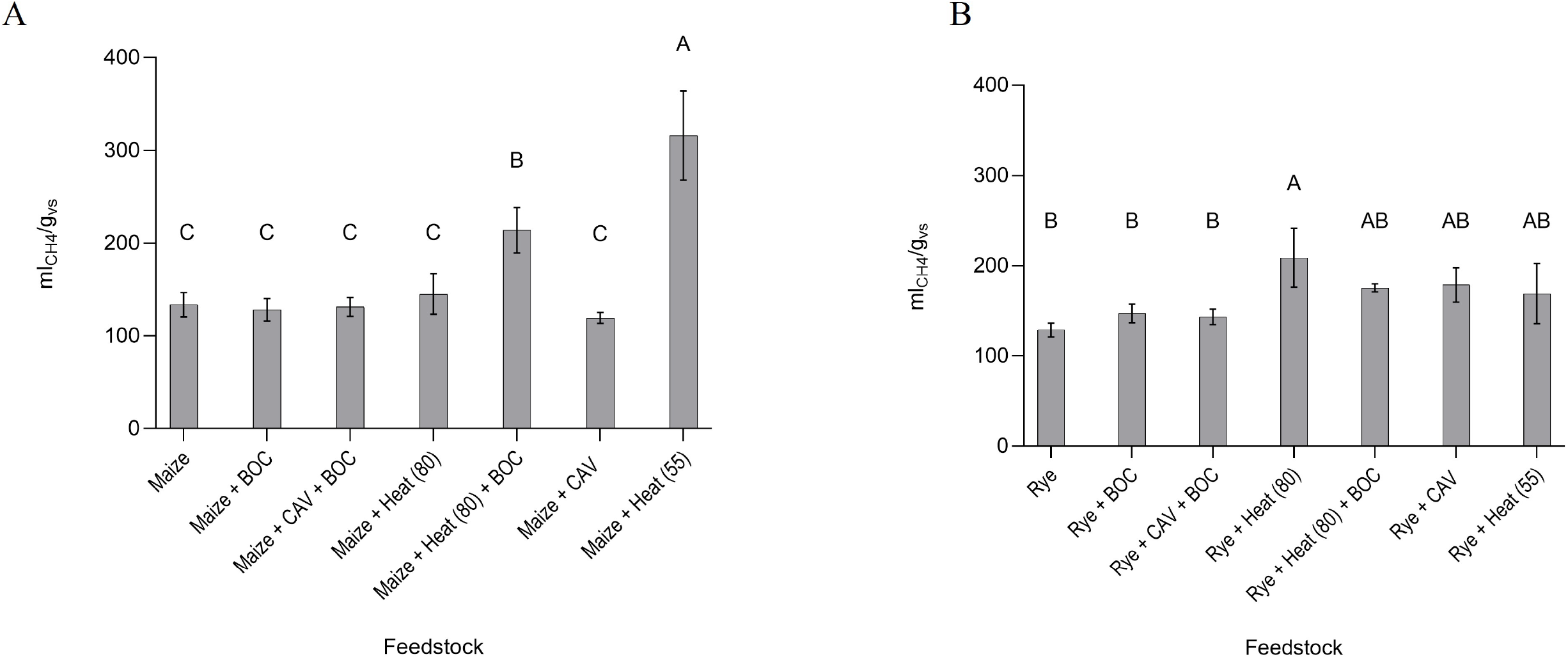
End time point of biomethane (ml CH_4_/g_vs_) production curve (cumulative) for two pretreated agricultural feedstocks. A) Maize, B) Rye. Both graphs have been subject to a one-way ANOVA with *post-hoc* Tukey tests where *p =<0.0001* (maize) and *p =0.0043* (rye). Full details of ANOVA results can be found in Supp. Table 2. All data presented as mean ±standard deviationof triplicate measurements. Results of Tukey’s HSD test of significant differences are shown on the graphs by letter.

## 4 Discussion

This study determined the effect of different pretreatments prior to AD on maize and rye, two feedstocks widely used in commercial biogas production. The aims of the project were to:

1. understand whether different pretreatments could influence the amount of organic material available for hydrolysis to increase biomethane yields
2. determine the success of the pretreatment methods relative to each other and their impact on different feedstocks for future use in commercial scale AD plants
3. Contextualise the results regarding implications for future development of the AD industry, particularly in the context of moving away from its current reliance on subsidies.

### 4.1 The overall effect of novel pretreatments on agricultural feedstocks

The maize and rye feedstock samples were pretreated at the ALPS testing plant facility (Chedglow, UK) using a range of mechanical, thermal, and biochemical methods. When compared to the control treatments, nearly all pre-treated samples exhibited an uplift in methane yield. As a current technological and economical bottleneck in AD, the need for quick, cheap, and relatively simple pretreatment methods for these feedstocks are required to improve the process [27]. The efficacy of the individual methods and the process energy demands at commercial stage for this experiment are discussed below.

### 4.2 The effect of heat pretreatments on two agricultural feedstocks

Thermal pretreatments have been well documented as effective means of overcoming the breakdown of lignocellulosic feedstock [28-30]; . However, separation of lignin from cellulose and hemicellulose employing this treatment requires significant energy that can prove too costly when offset against the increase in biomethane yield. Research into less energetically expensive methods of thermal pretreatment have been investigated in recent years, with a focus on recycling surplus heat and energy from existing plants, such as combined heat and power (CHP) [31, 32]. Although this method takes longer than conventional torrefaction, there is greater economic viability of pre-treating solid feedstocks over a longer period at a lower temperature to achieve an increase in biogas yields.

Both thermal pretreatments (heating the feedstock/water mix to 55 °C and 80 °C) increased the solubility of the feedstock and enhanced biogas yield in the experiment for both maize and rye. The results revealed that low-temperature heat pretreatment was the most effective method compared to cavitation and addition of BOC. It significantly increased the amount of biomethane (8.6 – 136.6%). Specifically, maize heated to 55 °C (136.6%) and rye heated to 80 °C (62%) exhibited the greatest increases in biomethane yield. This is consistent with findings in existing literature [33, 34], although it is important to note that the study conducted by Rademacher was run at thermophilic conditions with rye, and not technically a pretreatment, although it generated an increase in biogas yields.

Despite rye having a lower absolute methane yield compared to maize (3.4% lower), it generated a higher daily gas output rate and a notably shorter lag phase when heated prior to digestion (Table 2). However, it required a temperature of 80 °C to achieve this uplift. In this context the energy impact of 80 °C heating becomes a consideration. At commercial scale where waste process heat is available, 80 °C may be economic otherwise other pretreatments may be more energy efficient, albeit with gas being released more slowly.

Rye performed best at high temperature or under mechanical degradation. This may be explained by the cell structure and/or lignocellulosic content [35] but is beyond the scope of this study and requires further investigation to elucidate the breakdown components of the recalcitrant material.

### 4.3 The effect of mechanical pretreatment on agricultural feedstocks

Where it is not preferable to heat feedstocks due to economic considerations, other pretreatments may be more appropriate, such as mechanical means including cavitation.

Although cavitation does not remove lignin, it does lead to a reduction in particle size, resulting in an increase in surface area [28]. This pretreatment was most effective when combined with a chemical pretreatment [36] in increasing biogas yields, but has also been demonstrated to increase yields when used independently, also [37].

The third best performing rye sample (208 versus 208.7 (mL/gVS)) was cavitation. This suggests that rye requires more intensive degradation techniques than maize to increase conversion and maximise gas potential. In this context the energy impact of 80 °C heating versus cavitation becomes a consideration. At commercial scale where waste process heat is available, heating to 80 °C may be economic otherwise cavitation may be more energy efficient, albeit with gas being released more slowly.

### 4.4 Bio-Organic Catalyst (BOC) produces variable biomethane production which is feedstock dependent in a batch system

The results of the study have demonstrated that when used as an independent pretreatment BOC inhibited methane production in maize when compared to the control, and only marginally increased Biochemical Methane Potential in the rye feedstock. BOC and cavitation pretreatments increase gas yield (11 – 39%) and combined cavitation and BOC lowered HRT in the lag phase by 68%. (Table 2) The BOC acts as surfactant (reducing the lignin bonds and enabling greater access to cellulose and hemicellulose with the cavitated feedstock) [38]. Typically in a commercial setting, this required dual stage fermentation to optimise the BOC effect. The catalytic effect in a two-stage system (when not combined with other pretreatments) is not yet fully understood and is subject to ongoing commercial research. In single stage AD, the BOC is effective in breaking down FOGs but appears to require a two-step system to achieve increased bioavailability for AD. However, this was not the case in this single stage batch fermentation experiment when combined with other pretreatments. With this single stage fermentation and pretreatment protocol, there was a possibility that BOC acted as a competitor to the microbial communities (in the maize in particular), potentially impacting the overall efficiency of the AD process, resulting in greater lag time. The interaction between BOC and microbial communities in the sludge merits further investigation. Understanding this dynamic is essential for determining the use of BOCs in conjunction with heat pretreatments in single stage fermentation, ensuring that they compliment rather than hinder microbial activity. Also essential are further investigations into the two-step process as a continuous process at the commercial scale.

### 4.5 The implications of pretreatments on the Advanced Anaerobic Digestion (AAD) process

Approximately 740 AD plants currently operate in the UK [39]. A 15% increase in efficiency of a single 500 kW plant would lead to an increase of 657,000 kWh, saving 127 t/yr CO2 [4], compared with power from gas plants. In the longer term, these efficiency gains would allow existing market demand to be met with smaller bioreactors [40]. This has the potential to alter the economics of the whole sector by lowering costs and reactor size, de-risking the technology, and enabling greater asset portability and flexibility, but requires further investigation at the commercial scale [41].

This study has identified the opportunity of a novel pretreatment in combination with different industrial processing may offer as a means of increasing the yields and influencing hydraulic retention times in commercial biogas production. With increasing competition for, and environmental concerns in using conventional feedstocks, the ability to increase the conversion efficiency from the existing supply chain is appealing [30].

The study demonstrated that the protocols vary with the type of feedstock used and consequently so does the economics of the methodology. For example, grasses such as Rye, have a lower agricultural value than maize [42]. Therefore, applying heat pretreatment at 80 °C may be commercially viable for this feedstock, particularly if waste heat is available within an existing plant to be utilised. It may also be the case commercially that feedstocks could be blended [40, 41], with some of the feedstock blend receiving pretreatment and others not.

The results of this study demonstrate that maize heated to 55 °C and cavitated rye (with added BOC) provided the optimum conditions for gas production. The approaches examined in this study clearly have the potential to optimise fuel production from ADD economically and at commercial scale, providing a realistic path to post government subsidies, a significant contribution to UK net zero and facilitating circular bioeconomies [16].

## 5. Conclusion

This study has demonstrated the potential suitability of lignocellulosic material for AD at commercial scale when using pretreatments which enable increased biogas yields. Specific pretreatments have different outcomes when employed on specific feedstocks, which therefore have the potential to create a blended approach to produce economically viable methods at commercial scale. However, there are challenges in providing the energy and equipment required for these treatments to be implemented at commercial scale. Further work is required to demonstrate that these low energy mechanical, thermal, and chemical treatments can be widely adopted by the AD industry, and therefore become established as sustainable practice without government subsidies.

## Acknowledgements

The authors would like to thank Nigel Holmes from Severn Trent Green Power Cassington AD Facility for providing the AD sludge inoculum used in the experiment. The authors would also like to thank Ken Hodgson for his help with the sample preparation at ALPS Ecoscience.

## Author contributions

**Nichola Austen**: Conceptualisation, methodology, visualisation, funding acquisition, investigation, formal analysis, writing – original draft, writing – review and editing, supervision, project administration, **Helen Theaker**: Resources, writing – original draft, writing – review and editing, **Nicholas Tenci**: Investigation, formal analysis, writing – review and editing, **Alan Beesley**: Methodology, funding acquisition, resources, writing – original draft, writing – review and editing, **Ian Thompson**: Conceptualisation, methodology, supervision, funding acquisition, writing – review and editing,

---

**Supplementary Table 1.** Pretreatment collection and measurements for experiment.

| Sample | Origin | Measurement | Temperature heated (°C) | Time treated (hrs) |
| --- | --- | --- | --- | --- |
| Maize | Alps test plant | g | - | - |
| Rye | Alps test plant | g | - | - |
| Maize + cavitation | Alps test plant | g/L | 80°C | 48 |
| Rye + cavitation | Alps test plant | g/L | 80°C | 48 |
| Maize + BOC | Oxford laboratory | g/μl | - | 48 |
| Rye + BOC | Oxford laboratory | g/μl | - | 48 |
| Maize + cavitation + BOC | Alps test plant | g/L | 80°C | 48 |
| Rye + cavitation + BOC | Alps test plant | g/L | 80°C | 48 |
| Maize + heat (80°C) | Oxford laboratory | g/L | 80°C | 48 |
| Rye + heat (80°C) | Oxford laboratory | g/L | 80°C | 48 |
| Maize + heat (80°C) + BOC | Oxford laboratory | g/L/μl | 80°C | 48 |
| Rye + heat (80°C) + BOC | Oxford laboratory | g/L/μl | 80°C | 48 |
| Maize + heat (55°C) | Oxford laboratory | g/L | 55°C | 48 |
| Rye + heat (55°C) | Oxford laboratory | g/L | 55°C | 48 |

**Supplementary Table 2.** 1-way ANOVA results of SCOD & biomethane results from two agricultural feedstocks.

| Variable/Figure | Analysis | Transform | Treatment,<br>error DF | F | *P* value |
| --- | --- | --- | --- | --- | --- |
| SCOD, maize 1a | 1-way ANOVA – pretreatment | no | 6, 14 | 31.8 | <0.000001 |
| SCOD, rye 1b | 1-way ANOVA – pretreatment | no | 6, 14 | 6.368 | 0.002105 |
| Biomethane, maize 4a | 1-way ANOVA – pretreatment | no | 6, 13 | 28.4 | <0.0001 |
| Biomethane, rye 4b | 1-way ANOVA - pretreatment | no | 6, 11 | 6.333 | -0.0043 |

